# Endothelial specific reduction in Arf6 impairs insulin-stimulated vasodilation and skeletal muscle blood flow resulting in systemic insulin resistance

**DOI:** 10.1101/2023.05.02.539173

**Authors:** Md Torikul Islam, Jinjin Cai, Shanena Allen, Denisse G Moreno, Samuel I Bloom, R Colton Bramwell, Jonathan Mitton, Andrew G Horn, Weiquan Zhu, Anthony J Donato, William L Holland, Lisa A Lesniewski

## Abstract

**Background:** Much of what we know about insulin resistance is based on studies from metabolically active tissues such as liver, adipose tissue, and skeletal muscle. Emerging evidence suggests that the vascular endothelium plays a crucial role in systemic insulin resistance, however, the underlying mechanisms remain incompletely understood. ADP ribosylation factor 6 (Arf6) is a small GTPase that plays a critical role in endothelial cell (EC) function. Here, we tested the hypothesis that the deletion of endothelial Arf6 will result in systemic insulin resistance.

**Methods:** We used mouse models of constitutive EC-specific Arf6 deletion (Arf6^f/-^ Tie2Cre) and tamoxifen inducible Arf6 knockout (Arf6^f/f^ Cdh5Cre). Endothelium-dependent vasodilation was assessed using pressure myography. Metabolic function was assessed using a battery of metabolic assessments including glucose- and insulin-tolerance tests and hyperinsulinemic-euglycemic clamps. A fluorescence microsphere-based technique was used to measure tissue blood flow. Intravital microscopy was used to assess skeletal muscle capillary density.

**Results:** Endothelial Arf6 deletion impaired insulin-stimulated vasodilation in white adipose tissue (WAT) and skeletal muscle feed arteries. The impairment in vasodilation was primarily due to attenuated insulin-stimulated nitric oxide (NO) bioavailability but independent of altered acetylcholine- or sodium nitroprusside-mediated vasodilation. In vitro Arf6 inhibition resulted in suppressed insulin stimulated phosphorylation of Akt and endothelial NO synthase. Endothelial cell-specific deletion of Arf6 also resulted in systematic insulin resistance in normal chow fed mice and glucose intolerance in high fat diet fed obese mice. The underlying mechanisms of glucose intolerance were reductions in insulin-stimulated blood flow and glucose uptake in the skeletal muscle and were independent of changes in capillary density or vascular permeability.

**Conclusion:** Results from this study support the conclusion that endothelial Arf6 signaling is essential for maintaining insulin sensitivity. Reduced expression of endothelial Arf6 impairs insulin-mediated vasodilation and results in systemic insulin resistance. These results have therapeutic implications for diseases that are associated with endothelial cell dysfunction and insulin resistance such as diabetes.

## Introduction

Insulin resistance is a major driver of metabolic dysfunction and diabetes^1-3^. Indeed, reduced insulin sensitivity is one of the early events in the sequelae of the development overt diabetes^1-3^. Therefore, understanding the origin and mechanisms of insulin resistance is imperative to discover novel therapeutics for metabolic diseases such as diabetes^1,2^. Much of what has been described about insulin resistance in the literature suggests the underlying dysfunction originates from metabolically active tissues such as adipose tissue, skeletal muscle, and liver^1-4^. One such mechanism of insulin resistance that has been described in both obesity and advancing age is adipose tissue dysfunction^2^. Adipose tissue dysfunction in these settings leads to the accumulation of lipids in ectopic sites such as skeletal muscle and liver and this underlies peripheral insulin resistance^2^. Beyond this traditional view, emerging evidence suggests that endothelial cell dysfunction may play a critical role in the development of insulin resistance^5-8^. In healthy state, when blood glucose is high, insulin causes dilation primarily in the skeletal muscle microvasculature resulting in increases in blood flow and glucose uptake^7,9^.

However, in pathophysiological states such as occurs in diabetes, obesity, and aging, insulin-induced increase in skeletal muscle blood flow is compromised, contributing to insulin resistance^10-12^. Despite evidence that endothelial dysfunction and attenuated blood flow are concomitant with insulin resistance, the cellular and molecular mechanisms remain incompletely understood.

Recent discoveries identify a host of metabolic or transcriptional regulatory genes in the endothelium that play critical roles in endothelial and metabolic dysfunction^5,6^. For example, endothelial specific deletion of insulin receptor or insulin receptor substrates 2 impairs skeletal muscle glucose uptake^13,14^. Evidence exists that endothelial cell specific deletion of the transcription factor Forkhead Box O1 (FoxO1) increases skeletal muscle vascular density and suppresses obesity-associated metabolic dysfunction^15^. Moreover, manipulation of endothelial genes that influence inflammation also impact systemic metabolic function^8^. Notably, endothelial cell specific inactivation of NF-κB, a transcription factor that promotes inflammatory gene expression, prevents high fat diet induced metabolic dysfunction^16^. Together, these studies suggest that endothelial specific inflammatory mediators can be a target for arterial dysfunction as well as associated metabolic dysfunction. In a series of studies in the last decade, ADP ribosylation factor 6 (Arf6) has been described as an inflammatory mediator in the vascular endothelium, although its role in arterial and metabolic function remains largely unknown.

Arf6 is a highly conserved membrane bound small GTPase of the Ras superfamily that acts as a molecular switch which is active when bound to GTP and inactive when bound to GDP^17,18^. The activity of Arf6 is regulated by guanine exchange factors and GTPase activating proteins (i.e., GIT1)^17,18^. Hyperactivation of Arf6 has been reported in multiple human diseases and animal models of human disease, including cancer metastasis, sepsis, diabetic retinopathy and neuroinflammatory diseases^17-20^. The known role of Arf6 in critical endothelial functions and inflammatory signaling suggests that it may be a therapeutic target for many diseases. In the endothelium, Arf6 modulates non-canonical, NF-kB independent, inflammatory signaling^18^. In response to the inflammatory cytokine interleukin-1β, Arf6 promotes the disruption of vascular endothelial cadherin (VECAD) surface localization by endocytosis and enhances endothelial cell permeability^18^. Inhibition of Arf6 using siRNA prevented VECAD disruption and endothelial permeability^18^. These data demonstrate that Arf6 signaling is an inflammatory modulator of endothelial cell function. Since endothelial inflammatory signaling pathways are known to regulate systemic metabolic function, it is plausible to think that endothelial Arf6 may play an important role in arterial and metabolic function.

In this present study, we sought to test the hypothesis that endothelial specific deletion of Arf6 will modulate arterial and metabolic function. To test this hypothesis, we utilized a mouse model of endothelial specific Arf6 deletion that was generated on a whole-body Arf6 heterozygous background (Arf6^f/-^ Tie2Cre). We found that the deletion of endothelial Arf6 impairs endothelium dependent dilation (EDD) to insulin and results in systemic metabolic dysfunction. The Tie2-Cre mouse model was created on a whole body Arf6 heterozygous background to increase the recombination efficiency, however, we observed that Arf6 heterozygosity results in a metabolic phenotype that is distinct from wildtype mice. Additionally, Tie2-Cre is known to influence other cell types (i.e., macrophage) other than vascular endothelium. Therefore, to reduce confounding effects we generated a tamoxifen-inducible endothelial cell specific Arf6 deletion mouse model by crossing Arf6^f/f^ mice with Chd5-Cre transgenic mice. To examine the role of endothelial Arf6 on arterial and metabolic function, we examined EDD to insulin and acetylcholine in skeletal muscle and white adipose tissue feed arteries. To further examine the consequences of this attenuated insulin stimulated vasodilation, we performed a battery of metabolic assessments, as well as assessed skeletal muscle blood flow and glucose uptake in response to insulin.

## Methods

### Animal Model

In this study, we utilized functionally wildtype (Arf6^f/+^) and constitutive endothelial Arf6 deletion with whole body Arf6 heterozygous (Arf6^f/-^ Tie2 Cre+) mice that were developed and described by Dr. Weiquan Zhu^19^. These mice were maintained in the University of Utah Animal Facility in the standard shoe box cages on a 12:12 light: dark cycle with water and food ad libitum. We also obtained Cdh5 Cre+ and Arf6^f/f^ transgenic mice from our collaborators Dr. Weiquan Zhu. We crossed Chd5 Cre+ mice with Arf6^f/f^ mice to generate tamoxifen inducible EC-specific Arf6 KO (Arf6^f/f^ Chd5 Cre+) mice and WT littermates (Arf6^f/f^ Chd5 Cre- or Arf6^+/+^ Chd5 Cre+). Mice were studied at 2-3 months of age. All animal studies were performed in compliance with the Guide for the Care and Use of Laboratory Animals (2011)^22^ and were approved by the University of Utah and Veteran’s Affairs Medical Center-Salt Lake City Institutional Animal Care and Use Committees.

### Metabolic testing

Metabolic assays were performed as previously described^23,24^. For glucose, and insulin tolerance tests (GTT, ITT), mice were fasted 5-6 h in the morning and baseline blood glucose was measured using Precision Xceed Pro Glucometer in blood collected via a tail nick. An additional ∼30 µL blood was collected for measurement of plasma insulin.

Mice were injected with either dextrose (2g/kg, i.p., GTT) or insulin (1U/kg, i.p., ITT) and blood glucose was measured at 15, 30, 60, 90, and 120 min after injection. During the GTT, 15 min after the glucose injection, ∼30 µl blood was collected to measure plasma insulin during GTT. Blood for plasma separation was collected into heparinized microvette® CB 300 (Sarstedt) and centrifuged at 7500g for 5 minutes at 4°C. Plasma was collected and stored at -80°C for subsequent analysis. Plasma insulin was measured using an enzyme-linked immunosorbent assay kit (Chrystal Chem) according to manufacturer’s protocol. To induce obesity-associated metabolic dysfunction, we fed a cohort of WT and iECKO mice a high fat diet containing 60% calories from fat, 20% calories from protein and 20% calories from carbohydrate for 8 weeks (Research Diets: D12492).

### Hyperinsulinemic euglycemic clamp

Hyperinsulinemic-euglycemic clamps were performed as previously described^25^. Briefly, unrestrained mice were able to move freely while being continuously infused with insulin (2 mU/kg/min) and a variable infusion of 50% dextrose to allow for steady-state blood glucose of approximately 120 mg/dL. Constant infusion of ^3^H-glucose throughout the experiment and for 90 minutes before the clamp allowed for the quantification of glucose infusion rate and uptake. At the end of a 2-hour clamp, ^14^C-2-deoxyglucose (13 μCi/mouse) was administered during steady-state conditions.

### Quantitative polymerase chain reactions

Assessment of gene expression was performed as previously described^23,26^. Total mRNA was extracted from frozen liver, gastrocnemius muscle, and perigonadal adipose tissue using RNeasy Mini Kit (Qiagen) according to the manufacturer’s protocol. cDNA was synthesized from 800 ng of total mRNA using QuantiTect Reverse Transcription Kit (Qiagen) according to the manufacturer’s protocol. Quantitative PCR was performed on 96-well plates using Sso Fast™ Eva Green R Supermix (Bio-Rad) with the Bio-Rad CFX™ Real-Time System. Expression of the genes were normalized to 18s and fold change was calculated using the 2^-ΔΔCt^ method. Primer sequences were as follows: *18s* F 5′-TAGAGGGACAAGTGGCGTTC-3′, *18s* R 5′- CGCTGAGCCAGTCAGTGT-3′; *Arf6* F 5′-ATGGGGAAGGTGCTATCCAAAATC-3′, *Arf6* R 5′- GCAGTCCACTACGAAGATGAGACC-3’.

### Primary lung endothelial cell isolation

Two weeks after the last dose of tamoxifen was administered, mice were euthanized via exsanguination via cardiac puncture while maintained under isoflurane anesthesia. Lungs were collected, minced using sterile scissors, digested by collagenase-I (Worthington Biochemical Co/ LS004194) and filtered through a 70µm strainer. Cell suspensions were incubated with PECAM conjugated dynabeads. Cells were seeded in 100mm cell culture dishes at 37° C in the incubator. When the cells were confluent, the cell suspensions were sorted using ICAM conjugated dynabeads. The sorted cells were plated into 6-well plate cell culture dishes. When cells were 80-90% confluent, cells were collected for protein extraction and western blotting.

### Western blotting

Protein lysates were prepared from primary lung endothelial cells and human umbilical vein endothelial cells using ice cold RIPA buffer (Sigma Aldrich) with proteases and phosphatase inhibitor cocktails (Thermo Fisher). Protein concentration was measured using BCA assay (Thermo Fisher) according to manufacturer’s protocol. Protein expression was measured by standard western blot procedures using anti-mouse primary antibodies against Arf6 (rabbit monoclonal; 1:1000; 19kDa; Cell Signaling), β-actin (mouse monoclonal; 1:1000; 40kDa; Abcam), total Akt (pan-Akt; 1:1000; 60 kDa; cell signaling), phosphorylated Akt (s-473 p-Akt; 1:1000; 60 kDa; cell signaling), phosphorylated eNOS; 1:1000; cell signaling). Goat Anti-Rabbit and Anti-Mouse IgG (H+L)-HRP Conjugate (Bio-Rad) was used as the secondary antibodies. Images were visualized and quantified using Bio-Rad ChemiDoc™ XRS+ with Image Lab™ Software.

### Ex-vivo vasodilatory function

To examine endothelium-dependent dilation (EDD), gastrocnemius muscle and perigonadal white adipose tissue feed arteries were dissected from euthanized mice. Arteries were cleared of surrounding tissue and cannulated in the stage of pressure myograph (DMT Inc., Atlanta, GA, USA). Arteries were pre-constricted with 2 μM phenylephrine, EDD and the contribution of nitric oxide (NO) to dilation were measured in response to the cumulative addition of insulin (0.01 nM to 10 nM) or acetylcholine (10^−9^ to 10^−4^ M) in the presence and absence of the NO synthase inhibitor, L-NAME mmol/L, 30 min), as described previously^27,28^. NO bioavailability was assessed by subtracting insulin-induced maximal dilation in the presence of L-NAME from insulin-induced maximal dilation in the absence of L-NAME. Endothelium-independent dilation was assessed by measuring vasodilation in response to the cumulative addition of the exogenous NO donor sodium nitroprusside (SNP) (10^−10^ to 10^−4^ M)^28^. Vessel diameters were measured using MyoView software (DMT Inc., Atlanta, GA, USA). All dose-response data are presented as percent of possible dilation after preconstriction to phenylephrine. Arteries that failed to achieve ≥ 20% pre-constriction were discarded.

### Transfection and siRNAs

siRNA transfection was performed as previously described^19^.Briefly, siRNAs were diluted in 12.5% HiPerFectct Transfection Reagent (QIAGEN) in OptiMEM (Invitrogen, Thermo Fisher Scientific) and incubated for 10 to 20 minutes at room temperature. Passage 5-6 human umbilical vein endothelial cells were resuspended in EGM-2MV (Lonza) and combined with siRNAs, such that the final concentration of siRNA was 30 nM (all targets). Cells were plated and allowed to grow overnight, and then the growth media was replaced. Three days after the initial transfection, the cells were transfected a second time using the same HiPerFect/siRNA concentrations as above. Arf6 siRNA sequence: 5’- CAACGTGGAGACGGTGACTTA-3’ (QIAGEN SI02757286). QIAGEN All Stars Negative sequence was used as control.

### Measurement of blood flow

To quantify baseline and insulin-stimulated skeletal muscle blood flow, we utilized the fluorescent microsphere technique as previously described^29,30^. We performed the fluorometric assay 2 weeks after the last dose of tamoxifen.

### Surgical procedures for blood flow assessment

For microspheres and insulin infusion as well as reference blood sample collection, we performed surgical procedures as previously described^31^. Briefly, we inserted micro-Renathane MRE-025 catheter (Braintree Scientific, Inc.) via the right common carotid artery. The catheter was inserted through a small insertion made by 32 G needle. During the catheter insertion, we temporarily occluded the blood flow of the carotid artery at the side proximal to the heart. After we inserted the catheter, the occlusion was removed, and the catheter was advanced towards the heart and tied with 6-0 silk suture. To collect the reference blood sample, a second micro-Renathane MRE-010 catheter (Braintree Scientific, Inc.) was inserted via the saphenous artery^32,33^ in a similar fashion. We withdrew the reference blood at 10µL/min rate using an infusion/withdrawal pump (Harvard Apparatus).

### Microspheres infusion

To assess the baseline (5 hr. fasting) blood flow, we infused 1.5 × 10^5^ blue-green, 15µm diameter polystyrene microspheres (Thermo Fisher) via the carotid artery catheter while continuously collecting reference blood samples via the saphenous artery catheter. We next infused insulin (1 U/kg body mass) via the carotid artery catheter. 15 min after insulin infusion, we infused 1.5 x ^105^ red, 15µm diameter polystyrene microspheres via the carotid artery catheter while continuously collecting reference blood samples via the saphenous artery catheter. 5 min after the infusion of the red microspheres, mice were euthanized as previously described, skeletal muscle (gastrocnemius, soleus, and tibialis anterior) and diaphragm muscle were collected, weighed, and placed in 15 mL screw cap polypropylene conical tubes. The tissues were later digested using potassium hydroxide as previously described^29,30^.

### Calculation of Blood Flow

Digested tissue samples were placed in a 96-well plate and fluorescence intensity was measured using a plate reader. Samples were analyzed in triplicates and average values were used for blood flow quantification. The fluorescence intensity was normalized to reference blood samples and blood flow was reported as µL/min/g tissue mass. Tissue blood flow was calculated as follows^26.27^:

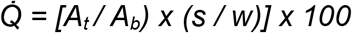

where 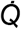 is blood flow (µL/min/g), *A*_*t*_ is the individual sample intensity, *A*_*b*_ is the reference blood sample intensity, *s* is the withdrawal rate (10 µL/min) of the reference blood sample, and *w* is the tissue mass (g).

### Assessment of capillary density

Skeletal muscle and mesenteric capillary density were measured using intravital microscopy as previously described^34^. Briefly, mice were anesthetized with isoflurane (2–3%) in 97-98% oxygen at 2 L/min flow rate and placed in the supine position on a heated platform (37 °C). Subsequently, the skeletal muscles were exposed, or intestines were mobilized, gently exteriorized, and placed into a warm isotonic 0.9% saline bath (37 °C). Intravital microscopy was performed with a CapiScope handheld video capillary microscope (KK Technology, Honiton, UK) to view the skeletal muscle or mesenteric microcirculation. The intravital microscope uses a sidestream dark field (SDF) camera that uses green light–emitting diodes. The green light is primarily absorbed by hemoglobin in RBCs in the microcirculation that allows RBCs to be viewed in contrast to the background. Microvascular density was analyzed using an automated capture and analysis system (GlycoCheck, MicroVascular Health Solutions LLC, Alpine, UT).

### Vascular permeability assay

The skeletal muscle vascular permeability assay was performed as previously described^20^. Briefly, 2 weeks after the induction of endothelial Arf6 deletion, WT and iECKO mice were anesthetized with isoflurane (2–3%) in 97-98% oxygen at 2 L/min flow rate. 50 mL of 60 mg/mL Evans blue solution (Millipore Sigma) in UltraSaline A was injected into the retroorbital sinus. After 5 h, skeletal muscle was collected and weighed. The dye was eluted in 1 mL formamide for 18 h at 70°C. Absorbances at 620 nm and 740 nm were measured with the absorbance at 740 nm being subtracted out. Finally, the absorbances were normalized to the tissue mass.

### Statistics

Statistical analyses were performed using GraphPad Prism Software version 9.2.0. Most group differences were determined by two-way ANOVA with Tukey post-hoc tests or student’s *t* test. Group differences in the glucose-, and insulin-tolerance tests and insulin stimulated suppression of free fatty acids were assessed by repeated measures ANOVA. Statistical significance was set at p<0.05 for all analyses. Data are presented as mean ± SEM.

## Results

### Endothelial cell specific deletion of Arf6 impairs insulin-stimulated vasodilation in white adipose tissue feed arteries

To examine the role of endothelial Arf6 on arterial function, we utilized functionally wildtype (Arf6^f/+^) and endothelial specific constitutive Arf6 knockout on whole body Arf6 heterozygous (Arf6^f/-^ Tie2Cre+) mice. We first assessed the impact of endothelial Arf6 deletion on endothelium dependent dilation (EDD) in response to vasodilatory agonists insulin and acetylcholine (ACh) in the white adipose tissue (WAT) feed arteries. EDD to insulin was lower in WAT arteries from Arf6^f/-^ Tie2Cre+ compared to Arf6^f/+^ mice (Figures 1A, 1B; p=0.002). In the presence of nitric oxide (NO) synthase inhibitor, L-NAME, dilation was lower in Arf6^f/+^ mice (Figures 1A, 1B; p=0.003), but no difference was observed in Arf6^f/-^ Tie2Cre+ mice (Figures 1A, 1B; p=0.97), resulting in NO bioavailability that was markedly lower in Arf6^f/-^ Tie2Cre+ compared to Arf6^f/+^ mice (Figure 1C; p=0.04). We did not find any difference in ACh-induced vasodilation between the groups (Figure 1D, 1E; p=0.96). ACh-induced EDD was lower in both groups in the presence of L-NAME (p≤0.001), but no difference was found between the groups (Figures 1D, 1E; p=0.71). Additionally, dilation to inorganic NO donor sodium nitroprusside, SNP, was not different between the groups (Figures 1F; p=0.65). Taken together, these results demonstrate that endothelial specific deletion of Arf6 impairs insulin-, but not ACh-, stimulated EDD due to attenuated NO bioavailability.

**Figure 1:**
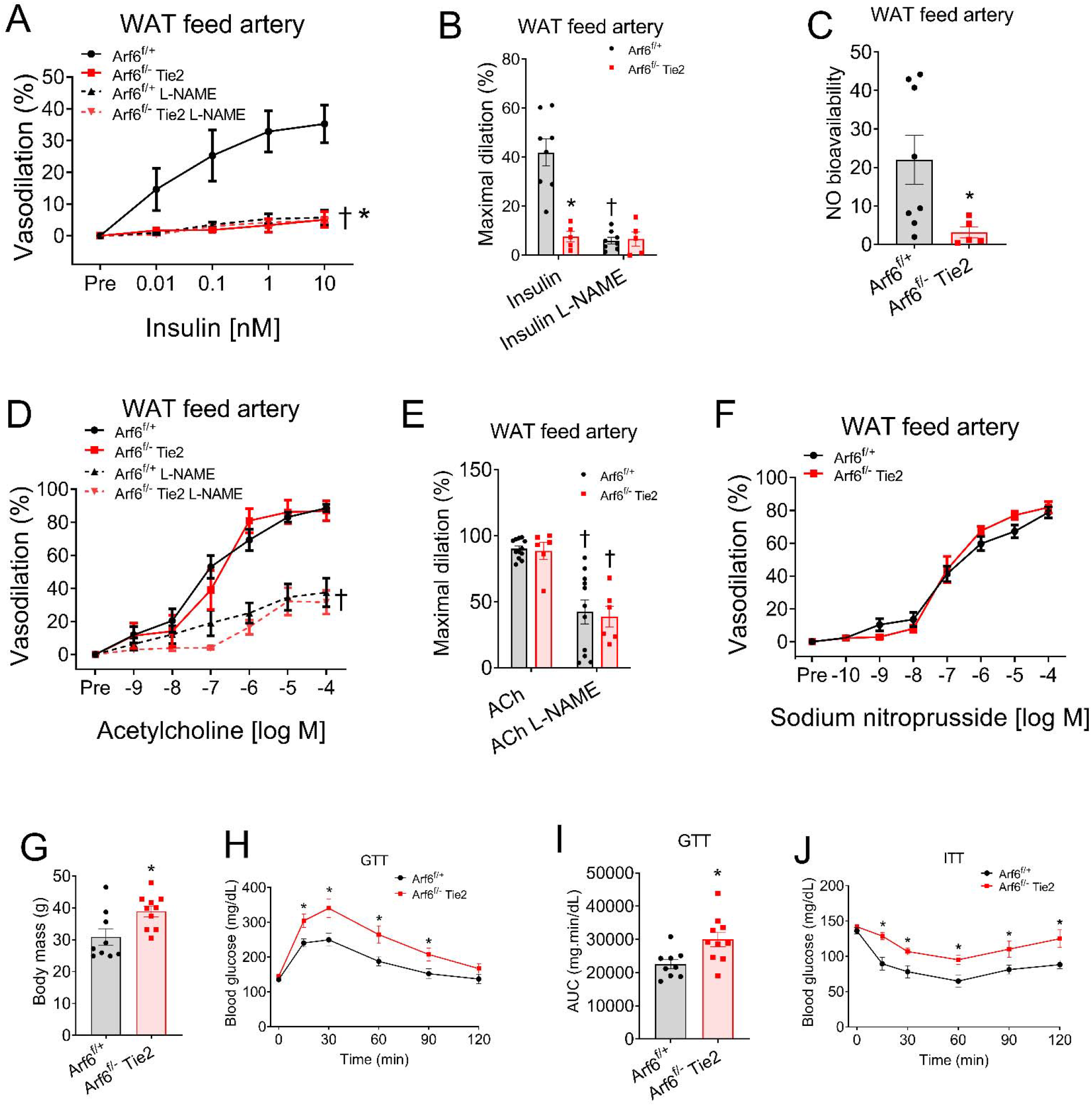
Endothelial Arf6 deletion impairs insulin-stimulated dilation in white adipose tissue feed artery and results in systemic metabolic dysfunction. (A, B) concentration response curves and maximal dilation for insulin in the absence- and presence- of nitric oxide (NO) synthase inhibitor, L-NAME from the white adipose tissue feed arteries of Arf6f/+ and Arf6f/- Tie2 Cre+ mice, (C) NO bioavailability for insulin from the white adipose tissue feed arteries, (D, E) concentration-response curves and maximal dilation for acetylcholine (ACh) in the absence- and presence- of nitric oxide (NO) synthase inhibitor, L-NAME from the white adipose tissue feed arteries, (F) oncentration-response curves for the endothelium independent vasodilator, sodium nitroprusside, (G) body mass, (H, I) blood glucose response curves and area under the curves (AUC) during a glucose tolerance test (GTT), (J) Blood glucose response curves during an insulin tolerance test (ITT). Data are shown as mean ± SEM with individual datapoints. N=6-10/group. Independent Student’s *t* tests, paired *t* tests and RM-ANOVA were performed to assess group differences. *p≤0.04 vs Arf6f/+; †p≤0.003 vs insulin or ACh without L-NAME.

### Impaired insulin-stimulated EDD is accompanied by systemic glucose intolerance and insulin resistance

We next assessed the effects of endothelial Arf6 deletion on metabolic function. The body mass of the Arf6^f/-^ Tie2Cre+ mice was higher than the Arf6^f/+^ mice (Figure 1G; p=0.01). Although we did not find a difference in baseline blood glucose (Figure 1H; p=0.15), blood glucose response curves and area under the curves during a glucose tolerance test (GTT) were higher in Arf6^f/-^ Tie2Cre+ compared to Arf6^f/+^ mice (Figures 1H, 1I; p≤0.01).

Likewise, the blood glucose response curve during an insulin tolerance test (ITT) was higher in Arf6^f/-^ Tie2Cre+ compared to Arf6^f/+^ mice (Figure 1J; p=0.008). Collectively, these results suggest that endothelial Arf6 deletion may result in systemic metabolic dysfunction.

### Generation and validation of tamoxifen inducible endothelial cell specific Arf6 knockout mouse model

While working with Arf6/Tie2 Cre mouse model, we observed that whole body Arf6 heterozygosity itself results in a metabolic phenotype that is distinct from wildtype or endothelial Arf6 knockout mice (data not shown here). Therefore, to investigate the direct contribution of endothelial Arf6, we developed a novel tamoxifen inducible endothelial specific Arf6 knockout (iECKO) mouse model by crossing the Arf6^f/f^ mice with Cdh5 Cre transgenic mice (Figure 2A). This model allowed for a significant reduction in endothelial Arf6 without the confound of whole body Arf6 heterozygosity. Mice were genotyped using polymerase chain reactions and agarose gel electrophoresis (Figure 2B). 2 weeks after tamoxifen administration (4mg/kg, 4 days, oral gavage), we found a ∼70% reduction (p=0.001) in Arf6 gene in EC-enriched carotid artery trizol effluents and a ∼30% reduction (p=0.02) in the lung, an endothelial-rich organ (Figure 2C). We did not find any difference in Arf6 gene expression in liver, skeletal muscle, white adipose tissue (WAT), or kidney (Figure 2C; p≥0.84), demonstrating the endothelial-specificity of our mouse model. To further validate the mouse model and examine the protein reduction efficiency, we measured Arf6 protein in primary lung endothelial cells isolated from iECKO and WT mice and found an 85-90% reduction in Arf6 protein in lung ECs from iECKO compared to WT group (Figures 2D, 2E; p≤0.001). Taken together, these data demonstrate that the administration of tamoxifen reduces both Arf6 gene and protein expression from endothelial cells but not from other organs of our novel mouse model.

**Figure 2:**
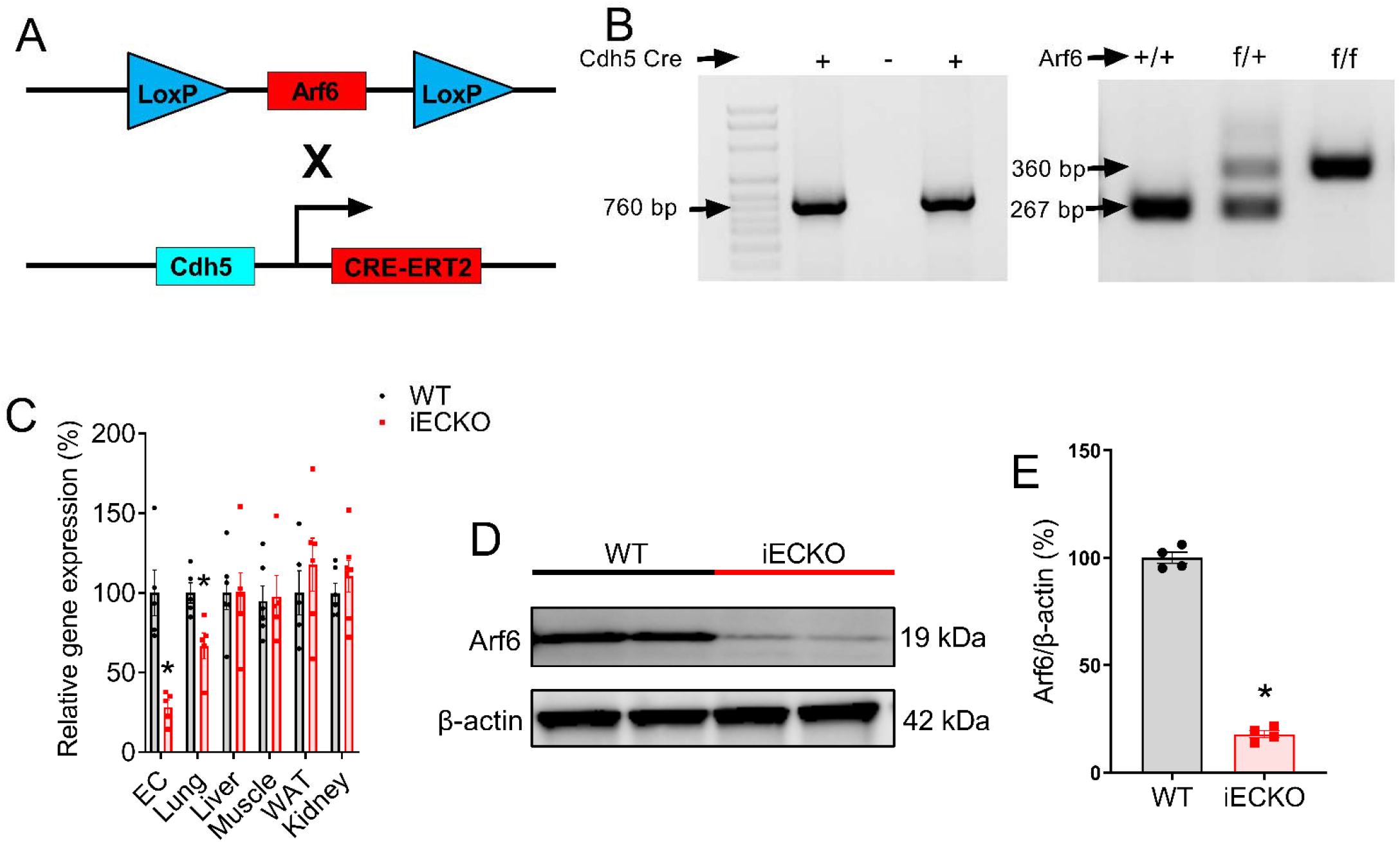
Generation and validation of tamoxifen inducible endothelial specific Arf6 deletion mouse model. (A, B) Generation and identification of endothelial cell (EC) specific tamoxifen inducible Arf6 knockout(iECKO) mouse model, (C) gene expression of Arf6 relative to 18S in EC-enriched carotid artery trizol effluents, lung, liver, muscle, white adipose tissue (WAT) and kidney 14 days after tamoxifen administration (4mg/day, 4 days, oral gavage), N=5-6/group, (D, E) representative western blotting images and densitometric quantification of Arf6 and β-actin proteins from primary lung endothelial cells (N=4/group. replicates were pooled from 3 mice) collected 14 days after tamoxifen administration. Data are shown as mean ± SEM with individual datapoints. Group difference was assessed using independent Student’s *t* test. *p≤0.02 vs WT.

### Induced reductions in endothelial Arf6 impair insulin-stimulated vasodilation in white adipose tissue and skeletal muscle feed arteries

After we generated and validated this new mouse model, we first sought to assess EDD to insulin and ACh of feed arteries from two key metabolic organs that contribute to insulin-stimulated glucose disposal, white adipose tissue, and skeletal muscle.

In white adipose tissue feed arteries, EDD to insulin was lower in iECKO mice compared to WT mice (Figures 3A, 3B; p=0.001). In the presence of L-NAME, dilation was lower in WT mice (Figures 3A, 3B; p≤0.001), but no difference was observed in iECKO mice (Figures 3A, 3B; p=0.19), resulting in NO bioavailability that was markedly lower in iECKO mice compared to WT mice (Figure 3C; p=0.002). However, EDD to ACh was not different between the groups (Figures 3D, 3E; p=0.66). In the presence of L-NAME, ACh-induced vasodilation was lower in both WT and iECKO mice (p≤0.001), although no difference was observed between the groups (Figures 3D, 3E; p=0.99). Additionally, dilation to inorganic NO donor sodium nitroprusside, SNP, was not different between the groups (Figure 3F; p=78).

**Figure 3:**
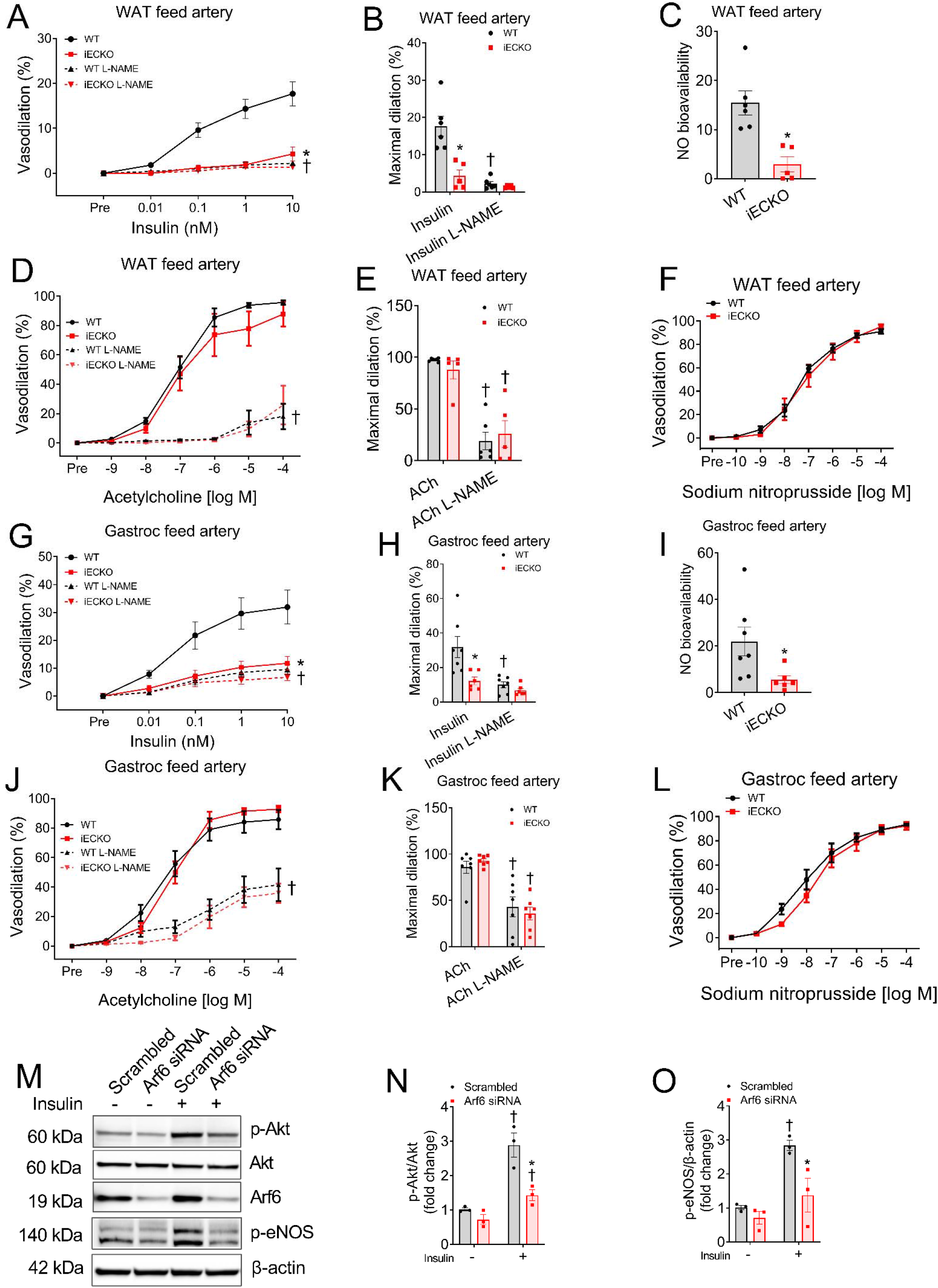
Endothelial cell specific reduction of Arf6 impairs insulin-stimulated dilation in white adipose tissue and gastrocnemius feed arteries. (A, B) Concentration response curves and maximal dilation for insulin in the absence- and presence- of nitric oxide (NO) synthase inhibitor, L-NAME from the white adipose tissue (WAT) feed arteries, (C) NO bioavailability for insulin from the WAT feed arteries, (D, E) concentration-response curves and maximal dilation for acetylcholine (ACh) in the absence- and presence- of nitric oxide (NO) synthase inhibitor, L-NAME from the WAT feed arteries, (F) concentration-response curves for the endothelium independent vasodilator, sodium nitroprusside from the WAT feed arteries, (G, H) concentration response curves and maximal dilation for insulin in the absence- and presence- of nitric oxide (NO) synthase inhibitor, L-NAME from the gastrocnemius muscle feed arteries, (I) NO bioavailability for insulin from the gastrocnemius muscle feed arteries, (J, K) concentration-response curves and maximal dilation for acetylcholine (ACh) in the absence- and presence- of nitric oxide (NO) synthase inhibitor, L-NAME from the gastrocnemius muscle feed arteries, (L) concentration-response curves for the endothelium independent vasodilator, sodium nitroprusside from the gastrocnemius muscle feed arteries, (M-O) representative western blotting images and densitometric quantification of p-Akt, Akt, p-eNOS and β-actin proteins from human umbilical vein endothelial cells (N=3/group). Data are shown as mean ± SEM with individual datapoints. N=5-7/group. Independent Student’s t test, paired t tests and RM-ANOVA were performed to assess group differences. *p≤0.03 vs WT or Scrambled; †p≤0.003 vs insulin or ACh without L-NAME or Ctrl.

EDD to insulin was lower in gastrocnemius feed arteries from iECKO compared to WT mice (Figures 3G, 3H; p=0.01). In the presence of L-NAME, dilation was lower in WT mice (Figures 3G, 3H; p=0.003), but no difference was observed in iECKO mice (Figures 3G, 3H; p=0.13), resulting in NO bioavailability that was markedly lower in iECKO mice compared to WT mice (Figure 3I; p=0.03). However, EDD to ACh was not different between the groups (Figures 3J, 3K; p=0.90). In the presence of L-NAME, ACh-induced vasodilation was lower in both WT and iECKO mice (Figures 3J, 3K; p≤0.001), although no difference was found between the groups (Figures 3J, 3K; p=0.40). Additionally, vasodilation to inorganic NO donor sodium nitroprusside, SNP, was not different between the groups. Taken together, these data are in agreement with the results from constitutive endothelial Arf6 knockout mouse model on the whole-body heterozygosity background.

### Reduction in endothelial cell Arf6 suppresses insulin-stimulated phosphorylation of Akt and eNOS

To assess the underlying mechanisms of impaired insulin-stimulated ex vivo vasodilation, we cultured human umbilical vein endothelial cells, knocked down Arf6 using small interfering RNA and measured baseline and insulin-stimulated phosphorylation of protein kinase B, commonly known as Akt, and endothelial NO synthase (eNOS). Reduction in Arf6 did not influence baseline phosphorylation of Akt and eNOS (Figures 3M-O; p≥0.11). However, Arf6 inhibition suppressed insulin-stimulated phosphorylation of Akt and eNOS (Figures 3M-O; p≤0.02). Taken together, these data suggest that attenuated eNOS phosphorylation and thus NO biosynthesis underlie the impairment in insulin-stimulated ex vivo vasodilation.

### Reduction in endothelial cell specific Arf6 results in systemic insulin resistance

To examine the impact of endothelial Arf6 reduction on systemic metabolic function, we performed a battery of metabolic assessments. We did not observe any difference in body mass between the groups either before or after tamoxifen treatment (Figure 4A; p≥0.71). Likewise, gastrocnemius muscle mass and white adipose tissue mass were not different between WT and iECKO mice (Figure 4B; p≥0.35). Baseline blood glucose and blood glucose response curves during a glucose tolerance test were not different between the groups (Figure 4C, p≥0.96).

**Figure 4:**
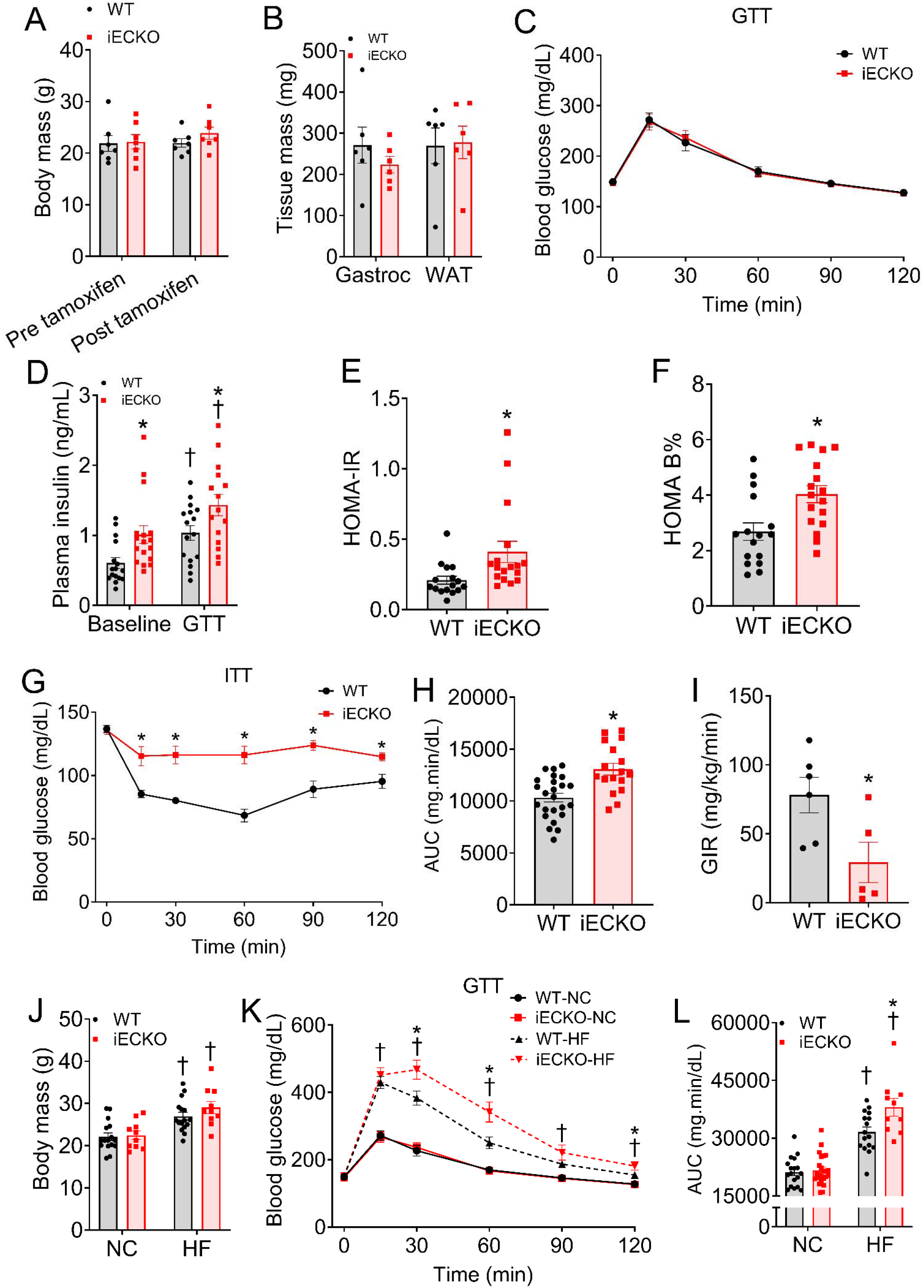
Endothelial cell specific reduction of Arf6 results in systemic insulin resistance. (A) Body mass of the wildtype (WT) and tamoxifen inducible endothelial Arf6 knockout (iECKO) mice before and after tamoxifen administration. (B) gastrocnemius muscle mass and white adipose tissue mass of WT and iECKO mice, (C) blood glucose response curves during a glucose tolerance test (GTT), (D) Plasma insulin level at baseline (6h fasting) and during GTT, (E) homeostatic model assessment for insulin resistance (HOMA-IR), (F) homeostatic model assessments for beta cell function (HOMA-B%), (G, H) blood glucose response curves and area under the curves during insulin tolerance test (ITT), (I) glucose infusion rate (GIR) during a hyperinsulinemic euglycemic clamp, (J) body mass of the WT and iECKO mice before and after the consumption of a high fat (HF) diet for 8 weeks, (K, L) blood glucose response curves and area under the curves (AUC) during glucose tolerance test of a normal chow (NC) and HF fed mice. Data are shown as mean ± SEM with individual datapoints. N=5-8/group for euglycemic clamps, body mass and tissue mass data; N=16-25/group for all other assays. Independent Student’s *t* test, paired *t* tests and RM-ANOVA were performed to assess group differences. *p≤0.03 vs WT; †p≤0.03 vs baseline or NC.

However, plasma insulin at baseline (6 hr fasting) was higher in iECKO compared to WT mice (Figure 4D; p=0.01). Plasma insulin during the GTT was higher in both iECKO and WT mice (Figure 4D; p≤0.03), however, iECKO mice exhibited higher plasma insulin compared WT (Figure 4D; p=0.01), suggesting insulin resistance. Likewise, homeostatic model assessments for insulin resistance and pancreatic beta cell function were higher in iECKO mice compared to WT (Figures 4E, 4F; p≤0.02). Blood glucose response curves and area under the curves during an insulin tolerance test were higher in iECKO compared to WT (Figure 4G, 4H; p≤0.001).

Likewise, glucose infusion rate during a hyperinsulinemic euglycemic clamp was lower in iECKO compared to WT mice (Figure 4I; p=0,03). Taken together, these results provide compelling evidence that an endothelial cell specific reduction of Arf6 results in systemic insulin resistance.

### Endothelial Arf6 reduction exacerbates obesity-associated glucose intolerance

Despite insulin resistance, the iECKO mice demonstrated a normal glucose tolerance. In the sequelae of type 2 diabetes, insulin resistance occurs at an earlier stage compared to glucose intolerance. Therefore, we next tested if endothelial Arf6 deletion exacerbates obesity-associated glucose intolerance. Consumption of a HF diet for 8 weeks increased body mass in both WT and iECKO mice (Figure 4J; both p≤0.0001); although no difference was found between the groups (Figure 4J; p≥0.15). HF diet-induced obesity resulted in elevated blood glucose response curves and area under the curves (AUC) during a glucose tolerance test (GTT) in both WT and iECKO mice (Figures 4K, 4L; p≤0.001). However, HF diet-fed iECKO demonstrated an exacerbated blood glucose response and AUC compared to HF diet-fed WT mice during GTT (Figures 4K, 4L; p≤0.01). Taken together, these data demonstrate that endothelial Arf6 reduction exacerbates the development of obesity-induced glucose intolerance.

### Systemic insulin resistance is concomitant with attenuated insulin stimulated blood flow and glucose uptake in the skeletal muscle

Tissue blood flow is a major contributor to peripheral insulin sensitivity. Therefore, we asked the question whether attenuated blood flow plays a role in insulin resistance resulting from a reduction in endothelial Arf6.

Baseline (6hr fasting) blood flow was lower in the skeletal muscle of iECKO mice compared to WT mice (Figures 5A, 5B; p=02). Stimulation with insulin (1 U/kg body mass, 15 min) significantly increased the skeletal muscle blood flow in WT mice (Figures 5A, 5B; p=0.007) but not in iECKO mice (Figures 5A, 5B; p=0.74). Likewise, skeletal muscle glucose uptake during a hyperinsulinemic euglycemic clamp was lower in iECKO compared to WT mice (Figure 5C; p=0.001). Baseline blood flow in the diaphragm muscle was not different between groups (Figures 5D, 5E; p=0.14). Stimulation with insulin significantly increased diaphragm muscle blood flow in WT mice (Figures 5D, 5E; p=0.04). Insulin-stimulated blood flow in the diaphragm muscle tended to increase in iECKO mice (Figures 5D, 5E; p=0.10). Diaphragm muscle glucose uptake during a hyperinsulinemic euglycemic clamp was not different between groups (Figure 5F; p=0.54), suggesting a differential impact of endothelial Arf6 reduction in different organs or tissues. Taken together, these data suggest that a consequence of attenuated EDD to insulin is reduced blood flow in the skeletal muscle resulting in impaired insulin-mediated glucose uptake and thus insulin resistance.

**Figure 5:**
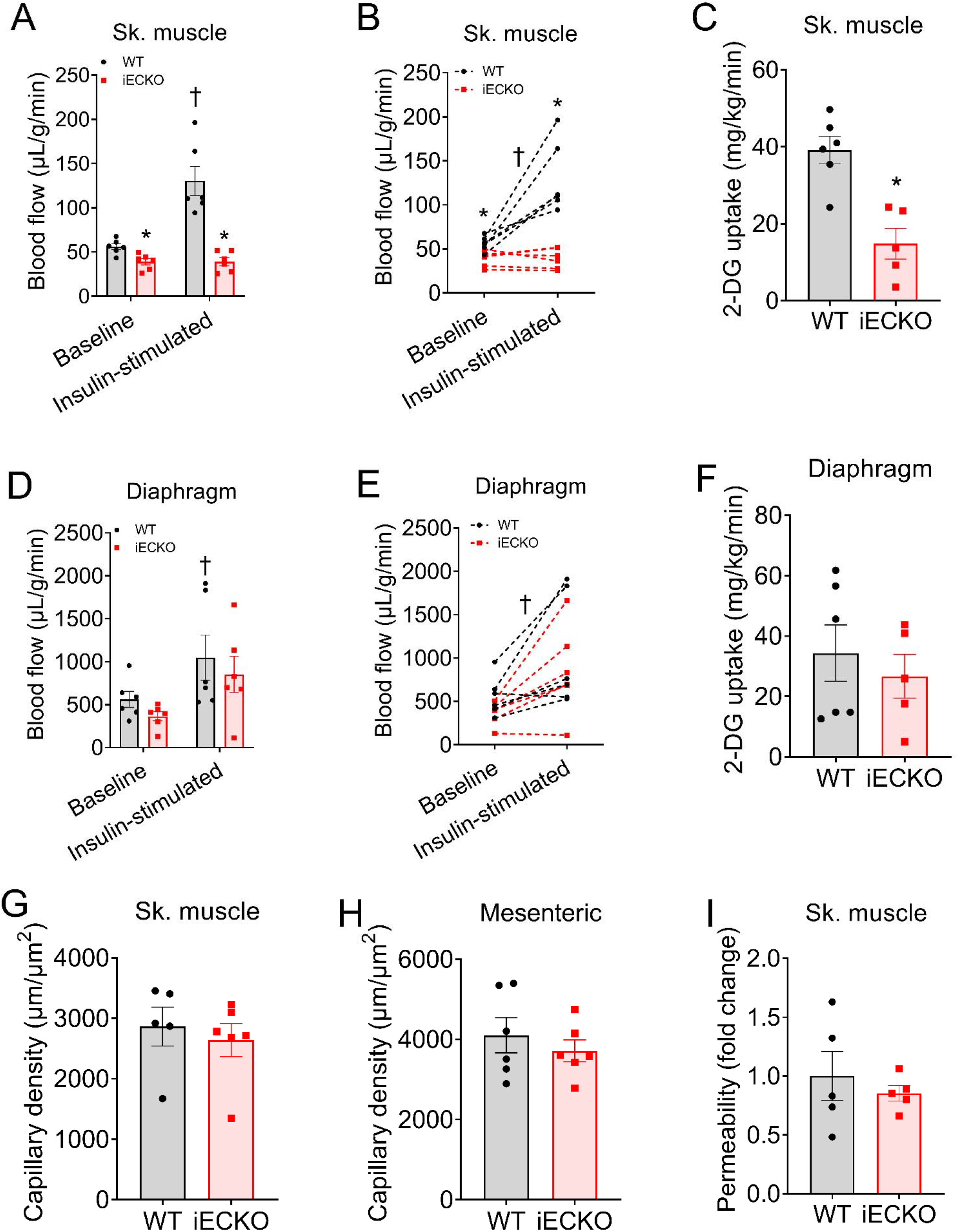
Endothelial Arf6 reduction leads to blunted insulin-stimulated blood flow response and glucose uptake in the skeletal muscle. (A, B) baseline and insulin stimulated skeletal muscle blood flow demonstrated as mean ± SEM with individual datapoints and paired response, (C) skeletal muscle glucose uptake during a hyperinsulinemic euglycemic clamp (D, E) baseline and insulin stimulated diaphragm blood flow demonstrated as mean ± SEM with individual datapoints and paired response, (F) diaphragm muscle glucose uptake during a hyperinsulinemic euglycemic clamp, (G) skeletal muscle and mesenteric capillary density, (I) skeletal muscle vascular permeability. Data are shown as mean ± SEM with individual datapoints. N=5-6/group. Independent Student’s *t* test and paired *t* tests were performed to assess group differences. *p≤0.02 vs WT; †p≤0.04 vs baseline.

### Endothelial Arf6 reduction did not alter capillary density and skeletal muscle vascular permeability

In addition to vasodilation, capillary density and vascular permeability may also influence blood flow, insulin sensitivity and glucose tolerance. Therefore, we examined the effects of endothelial Arf6 deletion on skeletal muscle and mesenteric capillary density as well as skeletal muscle vascular permeability. We did not find any difference in skeletal muscle or mesenteric arcade capillary density between WT and iECKO mice (Figures 5G, 5H; p≥0.47). Moreover, skeletal muscle vascular permeability did not differ between groups (Figures 5I; p=0.51). Taken together, these data suggest that the impact of endothelial Arf6 reduction on skeletal muscle blood flow is independent of capillary density and vascular permeability.

## Discussion

The purpose of this study was to examine the role of endothelial Arf6 in arterial function and subsequently, metabolic function. We made a novel observation that a reduction in Arf6 from endothelial cells impairs insulin-stimulated vasodilation in white adipose tissue and gastrocnemius muscle feed arteries due to reduced nitric oxide (NO) bioavailability. This genetic perturbation of endothelial cell specific Arf6 did not alter acetylcholine induced endothelium dependent or sodium nitroprusside-induced endothelium independent dilation in these arteries. Reduction in endothelial Arf6 suppressed insulin-stimulated phosphorylation of Akt and endothelial NO synthase (eNOS) and resulted in systemic insulin resistance due to attenuated insulin-stimulated skeletal muscle glucose uptake. Furthermore, endothelial Arf6 reduction resulted in exacerbated glucose intolerance in high fat diet fed obese mice. The insulin resistance and attenuated insulin stimulated glucose uptake was concomitant with reduced baseline and insulin stimulated blood flow in the skeletal muscle but was independent of capillary density or vascular permeability. Collectively, these data provide novel evidence demonstrating a critical role of endothelial Arf6 in insulin stimulated vasodilation, blood flow, and glucose uptake in skeletal muscle. These results advance our understanding of the vascular basis of insulin resistance and may have implications for designing novel therapeutics for diseases such as diabetes.

Endothelial cell function and systemic metabolic function demonstrate feed-forward interactions; however, the underlying mechanisms remain incompletely understood^8^. In this study, we provide evidence that a reduction in endothelial Arf6 impairs insulin-stimulated endothelium dependent dilation and that this is sufficient to induce systemic insulin resistance. The primary cause of attenuated insulin-stimulated vasodilation is reduced NO bioavailability. Mechanistically, Arf6 inhibition suppresses insulin-stimulated phosphorylation of Akt and eNOS and thus likely attenuates NO biosynthesis. The role of endothelial cell dysfunction and altered NO bioavailability on systemic metabolic dysfunction has been suggested by previous studies^5,6^. In a murine model of high fat diet-induced obesity, it has been observed that insulin-stimulated phosphorylation of eNOS decreases as early as one week after initiation of diet and that this precedes systemic metabolic dysfunction^35^. Indeed, evidence exists that insulin-stimulated vasodilation is critically important for skeletal muscle glucose uptake^36^. In fact, the vasodilatory effects of insulin on skeletal muscle arteries are blunted in states of insulin resistance such as obesity and advancing age^7,28^. Independent of vasodilation, NO may promote endothelial cell insulin uptake and transport through the barrier that is required for insulin stimulated glucose uptake^37^. Together, the role of NO and skeletal muscle artery vasodilation is well established as mediators of metabolism in the literature and our data demonstrate that Arf6 signaling is an important novel regulator in this interplay.

Genetic manipulation of several other signaling molecules in the vascular endothelium has also been demonstrated to influence systemic metabolic function. For example, endothelial cell specific deletion of insulin receptor substrate 2 (irs2) results in systemic insulin resistance via attenuated skeletal muscle glucose uptake^14^. Mechanistically, endothelial specific ablation of irs2 reduces insulin-stimulated phosphorylation of Akt and eNOS^14^. Evidence also exists that endothelial cell specific deletion of insulin receptor leads to insulin resistance partly because of delayed insulin-stimulated Akt phosphorylation in skeletal muscle and brown adipose tissue^13^.

Aside from these pathways, endothelial cell inflammatory signals also regulate systemic metabolic function. Notably, endothelial cell specific inactivation of a transcription factor, NF-kB, or tumor suppressor, p53 attenuate high-fat diet induced metabolic dysfunction^16,38^. Reduction in inflammatory signals prevents obesity-induced impairments in phosphorylation of Akt and eNOS^38,39^. Additionally, these genetic manipulations reduced the endothelial-derived secreted factors and thus dysfunction in metabolic tissues such as adipose and skeletal muscle^38,39^.

Taken together, these studies support that optimal endothelial cell function is critical to maintain systemic metabolic function and, here, we show that Arf6 signaling is essential to maintain endothelial and thus metabolic function.

A potential downstream consequence of impaired endothelium dependent dilation is a reduction in blood flow. In fact, it has been shown that obesity-related insulin resistance is associated with attenuated skeletal muscle blood flow^7,9,36,40^, suggesting a link between blood flow and insulin resistance. This possibility led us to test the hypothesis that impaired insulin-stimulated vasodilation and insulin resistance may accompany reduced insulin-stimulated skeletal muscle blood flow and glucose uptake. In support of our hypothesis, we observed that reducing endothelial Arf6 attenuates insulin-stimulated skeletal muscle blood flow and that this is concomitant with suppressed glucose uptake. In humans, it was found that insulin promotes leg blood flow in lean subjects in a NO dependent manner^41^. With obesity, insulin-stimulated blood flow is blunted^41^. In mice, deletion of endothelial specific irs2 reduced islet blood flow and insulin secretion, although an impact of this genetic manipulation on skeletal muscle blood flow was not tested^42^. It was also shown that endothelial irs2 deletion reduces skeletal muscle capillary blood volume, interstitial insulin level, and insulin stimulated glucose uptake^14^.

Collectively, evidence supports that organ perfusion and blood flow is a key macro mechanistic link for insulin action and, here, we provide evidence that endothelial Arf6 signaling is indispensable for this effect. While evidence exists that endothelial specific genetic perturbation of signaling molecules such as vascular endothelial growth factor (VEGF)^43^ or Forkhead Box O1 (FoxO1)^15^ improve systemic insulin sensitivity by increasing capillary density, endothelial Arf6 deletion did not influence capillary density or vascular permeability. These data suggest regulation of blood flow via dilation to insulin is a key component of insulin-stimulated glucose disposal and further that endothelial Arf6 is a mediator of this response.

## Conclusion, limitations, and future direction

Results from our study support the conclusion that endothelial cell Arf6 signaling is essential for insulin-stimulated endothelium dependent dilation of skeletal muscle and adipose tissue feed arteries. Consequently, endothelial Arf6 regulates insulin-stimulated blood flow and glucose uptake and thus systemic insulin sensitivity. Although, we demonstrate the role of insulin in regulating blood flow in our model, other mediators such as catecholamines may play a role in this regulation which requires future investigations. Although we show that Arf6 inhibition reduced the phosphorylation of Akt and eNOS, other signaling molecules such as MEK or PI3K may play important roles in this interplay which warrants future investigations. This study underlines the importance of vascular endothelium on systemic insulin sensitivity and identifies Arf6 as a critical regulator in this interplay. Currently, therapeutic targets for metabolic dysfunction are aimed at metabolically active tissues, such as adipose tissue, liver, and skeletal muscle. However, our results suggest that vascular endothelium may be a novel cell type that could be targeted to attenuate the burden of metabolic dysfunction and diseases.

## Author contributions

MTI and LAL were involved in the conceptualization, experimental design, and manuscript preparation. MTI, JC, SA, SIB, DG, RCB, AGH, JM, WLH, WZ, AJD and LAL were involved in data collection, analysis, or intellectual input. All authors read and approved the final version of this manuscript.

## Highlights

- Endothelial cell-specific deletion of Arf6 impairs insulin-stimulated vasodilation in adipose tissue and skeletal muscle feed arteries
- Arf6 inhibition suppresses the insulin-stimulated phosphorylation of Akt and eNOS in human umbilical vein endothelial cells
- Endothelial cell-specific Arf6 deletion results in insulin resistance in lean mice and glucose intolerance in obese mice
- The underlying mechanisms of insulin resistance is attenuated insulin-stimulated blood flow and glucose uptake in the skeletal muscle

